# FingerprintContacts: Predicting Alternative Conformations of Proteins from Coevolution

**DOI:** 10.1101/2020.04.13.037234

**Authors:** Jiangyan Feng, Diwakar Shukla

## Abstract

Proteins are dynamic molecules which perform diverse molecular functions by adopting different three-dimensional structures. Recent progress in residue-residue contacts prediction opens up new avenues for the *de novo* protein structure prediction from sequence information. However, it is still difficult to predict more than one conformation from residue-residue contacts alone. This is due to the inability to deconvolve the complex signals of residue-residue contacts, i.e. spatial contacts relevant for protein folding, conformational diversity, and ligand binding. Here, we introduce a machine learning based method, called FingerprintContacts, for extending the capabilities of residue-residue contacts. This algorithm leverages the features of residue-residue contacts, that is, (1) a single conformation outperforms the others in the structural prediction using all the top ranking residue-residue contacts as structural constraints, and (2) conformation specific contacts rank lower and constitute a small fraction of residue-residue contacts. We demonstrate the capabilities of FingerprintContacts on eight ligand binding proteins with varying conformational motions. Furthermore, FingerprintContacts identifies small clusters of residue-residue contacts which are preferentially located in the dynamically fluctuating regions. With the rapid growth in protein sequence information, we expect FingerprintContacts to be a powerful first step in structural understanding of protein functional mechanisms.

## Introduction

Proteins are dynamic entities that undergo significant conformational changes while performing diverse cellular functions such as drug binding,^1–4^ enzyme catalysis,^5–7^ and nutrient transport.^8–10^ Conformational changes often involve transitions among two or more alternative structures. Characterization of the conformational ensemble is critical for deciphering the relationship between protein structure and functional mechanisms.^11,12^ The past few years have seen substantial progress in characterizing protein conformations both experimentally and computationally.^13–15^ However, the prediction of protein conformational en-sembles still remains challenging. Experimental techniques such as X-ray crystallography or nuclear magnetic resonance (NMR) spectroscopy can only provide few snapshots from the conformational ensemble. In principle, molecular dynamics (MD) simulation is an attractive alternative due to its capability to model the behavior of proteins in full atomic detail.^16^ However, there are still several major challenges that limit its applicability. First, MD simulation requires at least one starting structure, but only 0.2% of known protein sequences^17^ have structures deposited in Protein Data Bank (PDB)18 as of 2018.^19^ Second, the accessible timescales in MD are still shorter than timescales for many functionally relevant conformational transitions. Third, the limited accuracy of force field models employed in simulations could impact the sampling of proper conformational ensemble.

Recently, rapid growth in sequence data has enabled us to exploit the evolutionary information embedded within multiple sequence alignments (MSAs) for computational prediction of native contacts in proteins. ^20–24^ As proteins evolve, they are subjected to evolutionary pressure to conserve their structures and functions, which can lead to covariance between residue pairs. These coevolving residue pairs are usually close in the native three-dimensional (3D) structure of the protein. The successful contact prediction from coevolving residue pairs has driven dramatic improvements in *de novo* protein structure prediction.^20–24^ According to Critical Assessment of Structure in Proteins (CASP),^25^ the flagship experiment of protein structure prediction, the average precision of contact prediction has increased from 22% in 2012 to 70% in 2018. The current methods for contact prediction can be divided into two groups: evolutionary coupling analysis (ECA) and supervised machine learning (SML).^26^ ECA methods rely on sequence information alone to identify the co-evolved residues as contacts, such as EVfold,^21^ PSICOV,^27^ CCMpred,^28^ and Gremlin.^29^ These ECA methods have been successfully applied to protein complexes,^30^ RNA structure prediction,^31^ and mutagenesis analysis.^32^ However, the quality of ECA methods highly depends on the quality of the multiple sequence alignments and it is required that the number of non-redundant sequences should be at least 64 times the square root of the length of the target protein.^33^ SML methods offer one promising avenue in these cases by integrating a variety of sequence-dependent and sequence-independent information. Popular SML methods include: SVMSEQ,^34^ CMAPpro,^35^ PconsC^2,36^ MetaPSICOV,^37^ PhyCMAP,^38^ CoinDCA-NN,^39^ and RaptorX-contact.^40^ SML methods have been found to be especially effective on proteins without many sequence homologs. ^41,42^ The increase in the accuracy of predicted residue-residue contacts has fueled the substantial improvements in *de novo* protein structure prediction.^43^

The question then becomes whether residue-residue contacts can be used to predict more than one conformation. Current techniques can only predict one single structure from residue-residue contacts due to the challenge of deconvoluting coevolutionary signals coming from different conformations. Considering the structural diversity of proteins, the coevolutionary signals contained in multiple sequence alignments could be a complex blend of different structural constraints imposed by the conformational ensemble.^44^ Recently, coevolutionary signals have been combined with physical models of proteins such as structure-based models (SBMs) to capture different conformational states. ^45–47^ These studies have shown that conformational diversity is embedded in coevolutionary signals. Unfortunately, the paradigm of integration of coevolutionary signals and protein physical models is applicable only when at least one conformation of the protein is available. In other words, current methods cannot identify more than one conformation from coevolution information alone.

Here, we propose a method called FingerprintContacts that combines residue-residue contacts and machine learning to quickly explore a small set of contacts that are relevant for conformational dynamics and predict alternative conformations of the protein by including or excluding these contacts as structural constraints. This algorithm was inspired by our previous work^48–50^ where we demonstrated that a small set of residue-residue contacts are sufficient to characterize complex conformational dynamics of proteins involved in folding and conformational change processes. However, pinpointing a few dynamically crucial contacts among the many evolutionarily coevolving residues without *a priori* knowledge remains a major challenge. To address this challenge, we hypothesized that the different selective pressures acting on a protein by the conformational ensemble are likely to result in a single conformation that dominates the structural prediction using all the top contacts. For example, we expect that the default structure, predicted using all the top contacts, will be structurally more similar to one conformer instead of being equally distant from two different alternative conformations. If the best conformer, the conformation that is the most similar to the default structure, exists, then it should be possible to identify another alternative conformation by removing a set of contacts maximizing the structural dissimilarity from the default structure while ensuring similar amount of contact satisfaction. Literature on protein sectors has shown that coevolving residues can be clustered into physically connected groups relevant for various biological functions, such as protein stability, enzymatic efficiency, or allostery.^51,52^

The FingerprintContacts leverages these ideas (1) to predict the default structure using all the top contacts as the best conformer, (2) to cluster all top scoring residue-residue contacts into small communities based on their physical positions, (3) to define a reward function which quantifies the potential of a newly predicted structure being an alternative conformation, (4) to explore few small communities of residue-residue contacts that maximize the reward function, and (5) to predict an alternative conformation by relaxing the structural constraints imposed by these small communities of residue-residue contacts. This approach differs from existing techniques for predicting protein conformational diversity using coevolutionary information in that it seeks to explore distant conformations of the protein by using the coevolution information alone rather than integrating it with structural information or dynamic simulation models. This is an important distinction because it offers two advantages: (1) widest scope, i.e. being able to obtain alternative conformations even for proteins without a PDB structure, and (2) computational efficiency, i.e. without tedious computational sampling of the enormous conformational space of protein. To test FingerprintContacts, we have applied it to eight ligand binding proteins due to the large conformational diversity and the availability of experimental structures for verification of our predictions. We begin by analyzing the default structure and contacts obtained from existing techniques to assess the two assumptions underlying the proposed algorithm. Then, we test FingerprintContacts’s ability to identify dynamically relevant residue-residue contacts and to predict alternative conformers.

## Methods

The main goal of this work was to develop a method that predicts distant conformations from residue-residue contacts alone. Our strategy is based on the analysis of residue-residue contacts to identify small clusters of contacts, fingerprint contacts, that cause significant structural perturbations in the default structure. Revealing such fingerprint contacts is critical because it may reflect the underlying dynamics involved in the conformational transition between two distant conformations. Furthermore, these fingerprint contacts can be excluded from structural constraints to predict an alternative conformation. Due to the hierarchical nature of protein evolution, a machine learning strategy particularly suited for this task is hierarchical clustering, which builds a hierarchy of clusters.^53^ Here we use agglomerative clustering to cluster residue-residue contacts based on their spatial closeness, which is a bottom-up approach, merging up pairs of clusters as moving up the hierarchy.^53^ The model parameters used for the eight test cases are listed in the SI (Table S1).

### FingerprintContacts Algorithm

The protocol developed, as outlined in Figure 1, is based on four consecutive stages. The algorithm parameters *n_clusters_range*, *bound*, *zscore_thres*, and *reward_thres* can be modified in the input file (see in SI for details). These parameters will affect the running time and the performance of the algorithm. The algorithm is implemented in the Python language and uses Numpy,^54^ Pandas,^55^ and Sklearn^56^ as dependencies. Due to the parallelizable nature, FingerprintContacts is considerably fast at predicting alternative conformations for the protein.

**Figure 1:**
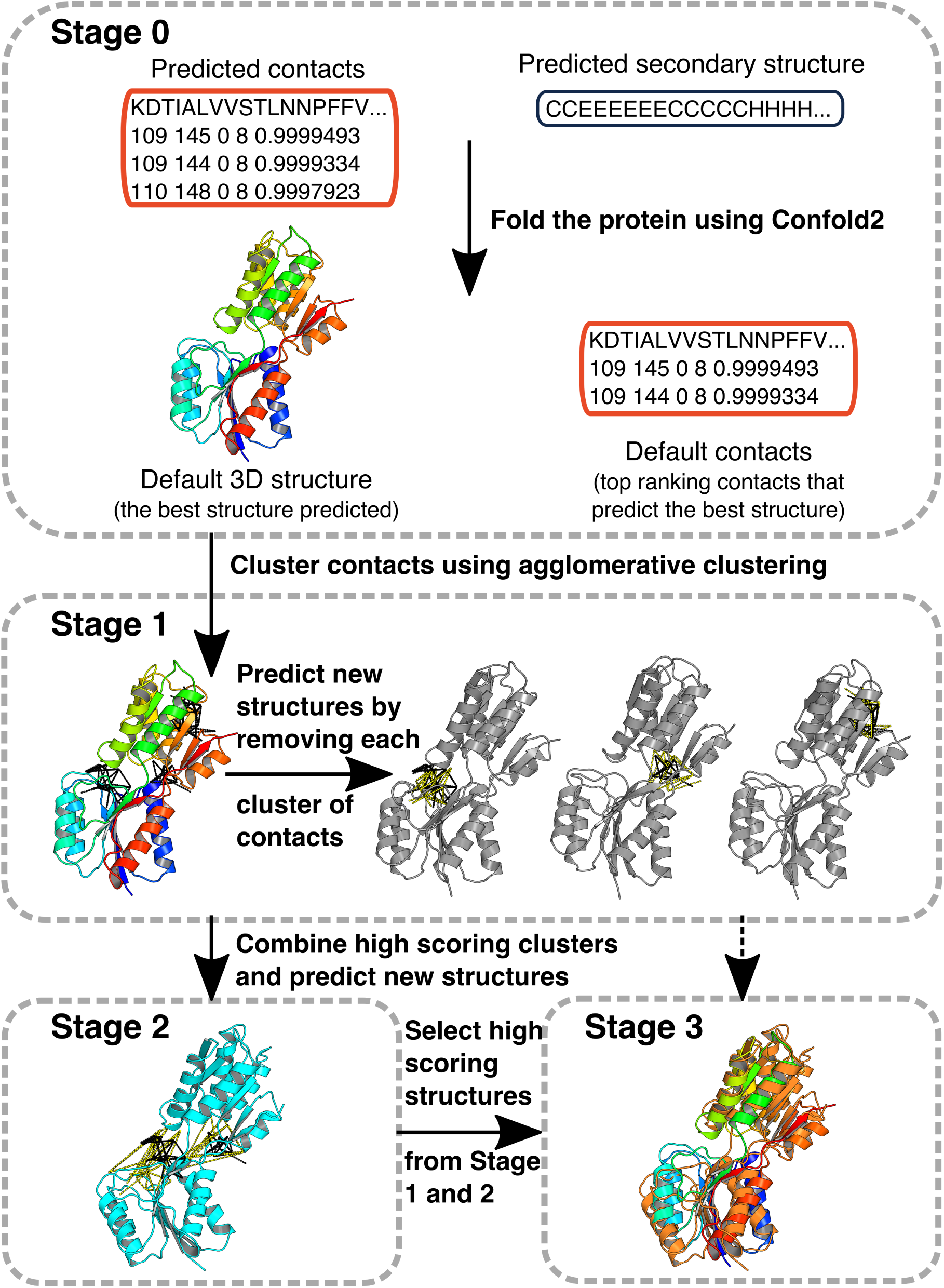
Overview of FingerprintContacts.

1. Identify the *Default 3D structure* (the best structure predicted) and *Default contacts* (the set of top ranking contacts that predicts the best structure). FingerprintContacts requires two inputs: predicted contacts and predicted secondary structure (see in Contact and Secondary Structure Prediction section for details). At stage 0, we feed the two inputs to Confold2,^57^ a powerful contact-driven *ab initio* protein structure prediction tool. The detailed description of Confold2 has been summarized elsewhere. ^57^ To determine the *Default contacts*, we build 20 models for each contact set consisting of top 0.1 L, 0.6 L,…, up to the top 4 L contacts (L: sequence length). The best model based on the satisfaction score of the top L/5 long-range contacts is then selected as the *Default 3D structure*, and the corresponding subset of contacts is selected as the *Default contacts*.
2. Cluster the *Default contacts* using agglomerative clustering. At stage 1, we cluster the *Default contacts* based on the Cartesian coordinates of the centers of the residue pairs using agglomerative clustering. To effectively explore all the small clusters, we perform agglomerative clustering with varied numbers of clusters. The algorithm parameter *n_clusters_range* specifies different numbers of clusters to use in the agglomerative clustering. For example, *n_clusters_range* = [10, 100, 10] provides 10 different number of clusters: 10, 20, 30,…, 100.
3. Select the small and non-redundant clusters. The previous agglomerative clustering generates a list of clusters which could be redundant due to the multiple clustering procedures performed using different numbers of clusters. We choose all the small and non-redundant clusters which satisfy the algorithm parameter *bound*. *bound* specifies the minimum and maximum size of a cluster. For example, *bound* = [0.005, 0.05, 0.1] constrains the size of a cluster in this stage to be no smaller than 0.005 of the size of *Default contacts* and be no larger than 0.05 of the size of *Default contacts*.
4. Predict new structures by removing structural constraints. For each small cluster obtained from the previous step, we generate a new contact file where the small cluster of contacts is removed from the *Default contacts*. We then feed the new contact file and the predicted secondary structure file to Confold2 in order to predict a new 3D structure. This results in a total of *M* new structures, where *M* is the number of small clusters.
5. Evaluate stage 1 structures and select candidate clusters. For each of the *M* clusters, we compute the reward function and zscore of rewards (see in Scoring Metric section for details). We choose clusters with *zscore_reward* greater than *zscore_thres* as candidate clusters for stage 2.
6. Generate new clusters by combining candidate clusters. At stage 2, we generate all the possible combinations of candidate clusters and select non-redundant combinations of clusters with size satisfying the algorithm parameter *bound*. As mentioned in step 3, *bound* constrains the size of clusters. For example, *bound* = [0.005, 0.05, 0.1] constrains the size of cluster in this stage to be no smaller than 0.005 of the size of *Default contacts* and be no larger than 0.1 of the size of *Default contacts*.
7. Predict new structure by removing structural constraints. Similarly to step 4, we predict *N* new structures by removing N different sets of contacts. *N* is the number of new clusters generated in step 6. This allows the algorithm to explore new structures by combining different communities of residue-residue contacts.
8. Evaluate stage 2 structures. For each of the *N* new clusters, we compute the reward function and zscore of rewards (see in Scoring Metric section for details).
9. Select high scoring structures from stage 1 and stage 2. At stage 3, we select high scoring structures with reward function greater than *reward_thres* from stage 1 and stage 2. It is important to note that we consider all the combinations among candidate clusters at step 6. This allows the algorithm to run multiple jobs in parallel to increase its efficiency. If the set of candidate clusters is very large or multiple processors are not available, this algorithm can be easily modified by growing the clusters in series. For example, we can combine 2 candidate clusters and then select high scoring combinations for the pairwise combination at the next step. This will dramatically decrease the total number of jobs.

### Clustering Metric

The clustering metric we selected for agglomerative clustering is based on the 3D position of center of each contact residue pair (c).

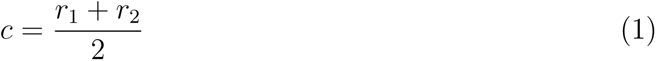

where *r*_1_ and *r*_2_ correspond to the Cartesian coordinates of residue 1 and residue 2, respectively.

### Scoring Metric

To measure how many *Default contacts* are satisfied in the given structure, we calculate the weighted contact satisfaction score (*Q*_*s*_):

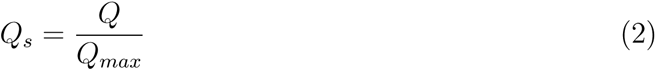

where *Q* is the sum of weights of satisfied contacts in the given structure and *Q*_*max*_ corresponds to the sum of weights of all the *Default contacts*. *Q*_*s*_ ranges from 0 to 1, from satisfying the least to the most number of contacts. The weights of residue-residue contacts are the probabilistic contact scores provided in the input contact file (see in Contact and Secondary Structure Prediction section for details). The weights are added to prefer the satisfaction of top ranking residue-residue contacts due to the rapid decay of true positive rate.

To measure the protein structural similarity, we computed template modeling score (*TMscore*) because it is independent of protein length and provides a clear quantitative cut-off (*TMscore* = 0.5) for protein fold definition.58 To be specific, proteins with *TMscore* ≥ 0.5 can be considered to have the same fold whereas proteins with *TMscore* < 0.5 do not have the same fold. TMscore is calculated using the equation:

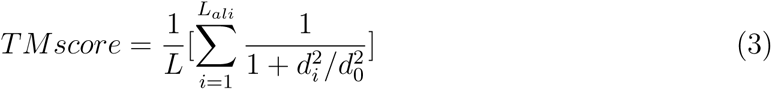

where *L* is the length of the target protein, *L*_*ali*_ is the number of shared residues in two proteins. *d*_*i*_ is the distance of *i^th^* pair of shared residues between two structures and *d*_0_ is defined to normalize *TMscore* to (0, 1]. The higher the *TMscore* is, the stronger the similarity is between two protein structures.

To quantify the relative likelihood of a new structure being an alternative conformation, we define the reward function by including both the contact satisfaction score *Q*_*s*_ and structural diversity score *TMscore* between the new structure and the *Default structure*. The *Q*_*s*_ component allows FingerprintContacts to ensure the new structure is correctly folded since previous studies suggested that the majority of residue-residue contacts are relevant for protein folding and are shared among different conformations. The *TMscore* component encourages FingerprintContacts to explore other conformations that are structurally different from the *Default structure*. The reward function is defined as:

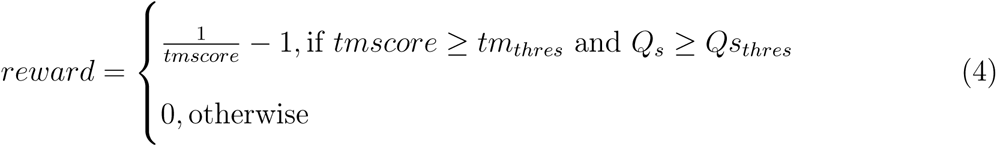

where *tmscore* is *TMscore* between the new structure and the *Default 3D structure*. *Q*_*s*_ is the weighted contact satisfaction score as defined above. The user-defined *tm*_*thres*_ should be no smaller than 0.5 since the different conformations of the same protein usually share the same fold. By default, we choose *tm*_*thres*_ = 0.5 to ensure *reward* ranges from 0 to 1.

To determine which clusters to select as candidates for the stage 2, we computed zscore of the reward, which measures the number of standard deviations a given point from the mean of all rewards.^59^ Using zscore facilitates the identification of significant clusters based on the distribution of the reward instead of explicitly specifying the number of clusters. In this work, *zscore*_*reward* is defined as:

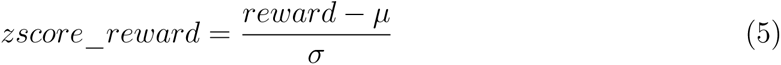

where *zscore*_*reward* is the zscore of reward, *reward* is the reward for a given cluster, *µ* is the mean of rewards, and *σ* is the standard deviation of rewards.

### Contact and Secondary Structure Prediction

We predicted contacts and secondary structures using the RaptorX-Contact web server.^40^ RaptorX-Contact applies an ultra-deep convolutional residual neural network to predict contacts and assigns each contact pair a weight, quantifying the possibility of the *Cβ − Cβ* distance (*Cα* for GLY) to be at or below 8 Å. We used these contact pair weights for the calculation of weighted contact satisfaction score *Q*_*s*_.

### Protein Data Set

To validate the FingerprintContacts algorithm, we tested its capability to predict known alternative conformations for eight proteins with varying conformational movements (Table 1).

**Table 1:** Protein used for benchmarking

| Uniprot Name | PDB-closed (chain) | PDB-open (chain) | Residues | Motion | TM |
| --- | --- | --- | --- | --- | --- |
| KAD_AQUAE | 3SR0 (B)<sup>60</sup> | 2RH5 (A)<sup>61</sup> | 206 | Concerted | 0.7100 |
| KAD_ECOLI | 1AKE (A)<sup>62</sup> | 4AKE (A)<sup>63</sup> | 214 | Concerted | 0.6841 |
| LIVK_ECOLI | 1USI (A)<sup>64</sup> | 1USG (A)<sup>64</sup> | 346 | Open-close | 0.6461 |
| RBSB_ECOLI | 2DRI (A)<sup>65</sup> | 1BA2 (A)<sup>66</sup> | 271 | Open-close | 0.6071 |
| AROA_ECOLI | 2AAY (A)<sup>67</sup> | 1EPS (A)<sup>68</sup> | 427 | Rotation-close | 0.6166 |
| ARGT_SALTY | 1LST (A)<sup>69</sup> | 2LAO (A)<sup>69</sup> | 238 | Open-close | 0.7044 |
| KAD4_HUMAN | 3NDP (B)<sup>70</sup> | 2AR7 (A)<sup>71</sup> | 231 | Rotation-close | 0.6794 |
| KAD2_YEAST | 2AKY (A)<sup>72</sup> | 1DVR (B)<sup>73</sup> | 220 | Concerted | 0.8224 |

## Results and Discussion

### Default structure strongly resembles the closed conformer

It is important to recognize that protein adopts an ensemble of conformers instead of a single unique structure.^74^ Many researchers have noted that residue-residue contacts are a complex blend of signals from the conformational ensemble.^23,45^ Therefore, we wondered whether the *default structure*, the structure that maximally satisfies the residue-residue contacts, is equally distant from different protein conformations. In other words, whether the conformational specific contacts from different conformers impact the final structure prediction to the same degree.

We calculated the TMscore, a measure of protein structural similarity, of the *default structure* against alternative protein structures (Table 2). Whereas the *default contacts* belong to both open and closed conformations (Table 3), *default structure* is structurally more similar to the closed than the open state of the protein (Table 2). We observed that TMscore in reference to the closed state is much higher than TMscore compared to the open state, with a difference around 0.17 on average (Table 2 and Table S2). For a closer examination of the results, we superimposed the *default structure* (green) and the open structure (magenta) onto the closed structure (blue) (Figure 2, Figure S1). We found that *default structure* is similar in overall structure to the known closed conformation in all eight cases but distinct from the known open structure.

**Table 2:**
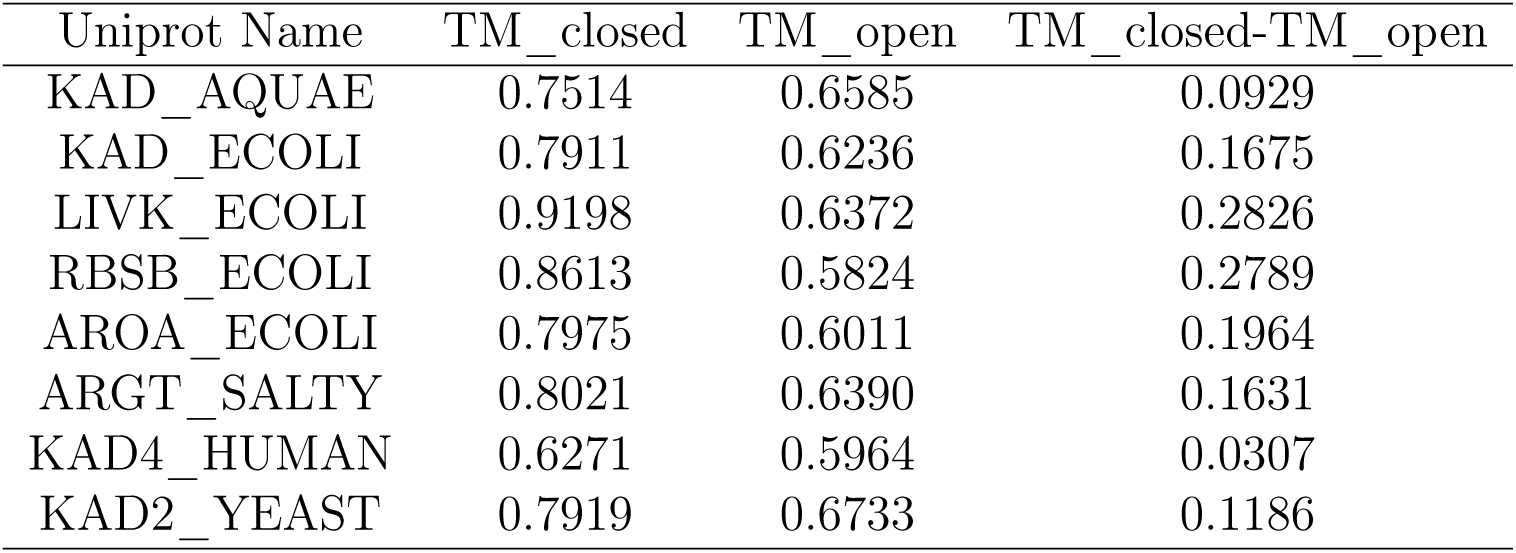
Comparison of *default structure* against both conformers of a protein

**Table 3:** Groups of residue-residue contacts

| Uniprot<br>Name | Optimal<br>number | $Q_s$ | | Conformation specific (%) | | Shared<br>contacts (%) | False<br>positives (%) |
| --- | --- | --- | --- | --- | --- | --- | --- |
|  |  | Closed | Open | Closed | Open |  |  |
| KAD_AQUAE | 330 | 0.7693 | 0.7767 | 6.67 | 6.67 | 68.18 | 18.48 |
| KAD_ECOLI | 235 | 0.8877 | 0.9068 | 3.40 | 5.53 | 84.68 | 6.38 |
| LIVK_ECOLI | 484 | 0.9480 | 0.9178 | 4.13 | 1.03 | 90.50 | 4.34 |
| RBSB_ECOLI | 650 | 0.8923 | 0.8693 | 4.77 | 2.46 | 82.92 | 9.85 |
| AROA_ECOLI | 640 | 0.8203 | 0.7993 | 3.91 | 1.72 | 77.66 | 16.72 |
| ARGT_SALTY | 262 | 0.9699 | 0.9553 | 2.29 | 0.76 | 94.66 | 2.29 |
| KAD4_HUMAN | 254 | 0.8051 | 0.8487 | 6.78 | 6.36 | 78.81 | 8.05 |
| KAD2_YEAST | 506 | 0.6940 | 0.6945 | 6.57 | 6.37 | 57.17 | 29.88 |

**Figure 2:**
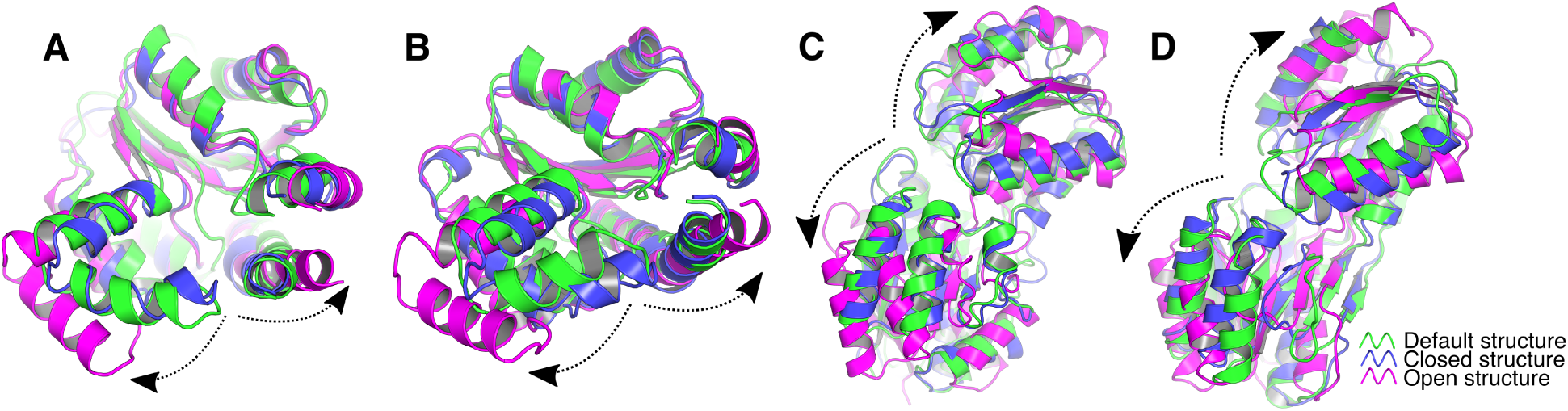
Structural superposition of default structure against both conformers of a protein. (A) KAD_AQUAE, (B) KAD_ECOLI, (C) LIVK_ECOLI, and (D) RBSB_ECOLI. Default structure, closed, and open structures are represented in green, blue, and magenta. The black lines represent the conformational motions.

It is established that the closed conformer is relevant for ligand binding and protein biological activities. Thus, it is possible that the closed conformation is under the strongest selective pressure during evolution. This explains why the structure predicted using all the top ranking residue-residue contacts, a mixture of signals from the conformational ensemble, is associated with the closed conformer. Taken together, this evidence supports the hypothesis that there exists a single conformation (closed conformation for the ligand binding protein) that dominates the structural constraints derived from multiple sequence alignments and consequently the prediction of *default structure*. Given that the *default structure* captures the closed conformation, we anticipate that the removal of the residue-residue contacts only consistent with the closed conformation from the folding calculation might help to identify the open conformation.

### Conformation specific contacts: Small but mighty

Due to the bias of the *default structure* towards the closed conformation, we next ask how many residue-residue contacts are conformation specific contacts. To this end, we classified the *default contacts* based on their *C*_*β*_ distances in the alternative conformations (Table 3, Figure 3, Figure S2). We considered a residue pair in contact if *C*_*β*_ distance ≤ 8 Å. We defined the following groups:

**Figure 3:**
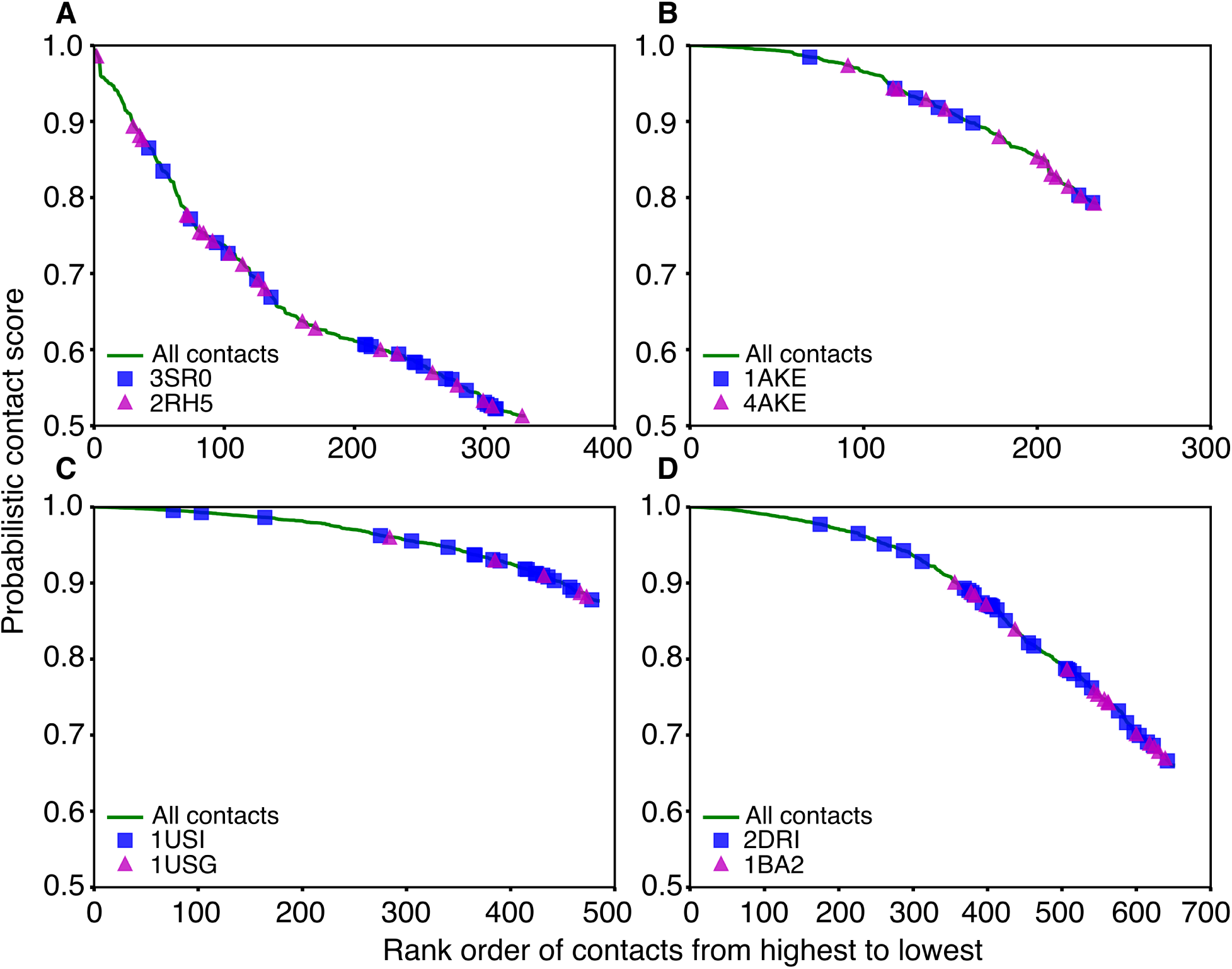
Distribution of conformation specific contacts in the alternative conformers of a protein. (A) KAD_AQUAE, (B) KAD_ECOLI, (C) LIVK_ECOLI, and (D) RBSB_ECOLI. All the *default contacts*, residue-residue contacts unique to closed and open structures are shown in green, blue, and magenta, respectively.

1. conformation specific contacts, that is, contacts that only exist in one of the conformers;
2. shared contacts, that is, contacts that exist in both of the alternative conformations; and
3. false positives, that is, contacts that do not exist in either of the conformations.

We observe that a large fraction of the residue-residue contacts (∼80% on average) are shared between the alternative conformers, which could be intra-protein contacts defining the fold of the protein family (Table 3). Second, a relatively small fraction of the residue-residue contacts (∼5% on average) are specific to a single conformer (Table 3). Third, conformation specific contacts rank lower among all the *default contacts*, which indicates that conformation specific residue pairs might coevolve less strongly than the shared residue pairs (Figure 3, Figure S2). This observation is consistent with a previous study which shows that the most strongly coupled residue pairs correspond to intra-protein contacts defining the fold of proteins. ^44,75^ Fourth, although the *default structure* is more similar to the closed conformation, the number of satisfied residue-residue contacts in closed structure is not relatively higher than those satisfied in the open structure (Table 3). On the contrary, the weighted contact satisfaction score (*Q*_*s*_) in the closed conformation is equivalent to *Q*_*s*_ in the open conformation (Figure S3). In the case of KAD_AQUAE, KAD_ECOLI, KAD4_HUMAN, and KAD2_YEAST, the weighted contact satisfaction score (*Q*_*s*_) in the closed conformation is actually smaller than *Q*_*s*_ in the open conformation. This indicates that the strength, instead of the number of conformation specific contacts, affects the final prediction results.

The above observations imply that there are two characteristics of conformation specific contacts: (1) a small fraction of the *default contacts* and (2) low ranking score. In this work, we utilize these features to explore the alternative open structure by identifying and excluding the fingerprint contacts, a small set of contacts that cause large structural deviations from the *default structure* without sacrificing the satisfaction of *default contacts*. To avoid the prediction of an incorrectly folded structure, we constrained the size of the set of removed contacts and added the weighted contact satisfaction score (*Q*_*s*_) to the reward function, which quantifies how many *default contacts* are realized in the new structure (as described in the Methods section).

### FingerprintContacts captures the alternative open state

Having demonstrated the two assumptions underlying our proposed method, that is, (1) the *default structure* resembles the closed conformation, and (2) the conformation specific contacts are a small fraction of the *default contacts*, we then analyzed the performance of FingerprintContacts on eight ligand binding proteins (Table 4, Table S3, Figure S4) which undergo large conformational changes when transitioning between open and closed state. Ligand binding proteins are a particularly interesting class of proteins because they can potentially be used in many medicinal applications, e.g. drug targeting and biosensing of disease markers. ^76^ Therefore, the prediction of alternative conformations could provide biophysical insight into the underlying mechanisms of ligand binding and create new therapeutic opportunities.

**Table 4:** A summary of FingerprintContacts results

| Uniprot Name | PDB-A<br>(chain) | PDB-B<br>(chain) | Blind top |  | Best |  |
| --- | --- | --- | --- | --- | --- | --- |
|  |  |  | TM_closed | TM_open | TM_closed | TM_open |
| KAD_AQUAE | 3SR0 (B)<sup>60</sup> | 2RH5 (A)<sup>61</sup> | 0.6303 | 0.7823 | 0.6702 | 0.8362 |
| KAD_ECOLI | 1AKE (A)<sup>62</sup> | 4AKE (A)<sup>63</sup> | 0.5548 | 0.6842 | 0.6445 | 0.7625 |
| LIVK_ECOLI | 1USI (A)<sup>64</sup> | 1USG (A)<sup>64</sup> | 0.5032 | 0.5907 | 0.5831 | 0.7361 |
| RBSB_ECOLI | 2DRI (A)<sup>65</sup> | 1BA2 (A)<sup>66</sup> | 0.5154 | 0.6502 | 0.5676 | 0.6961 |
| AROA_ECOLI | 2AAY (A)<sup>67</sup> | 1EPS (A)<sup>68</sup> | 0.4862 | 0.5921 | 0.5663 | 0.7757 |
| ARGT_SALTY | 1LST (A)<sup>69</sup> | 2LAO (A)<sup>69</sup> | 0.5060 | 0.5879 | 0.6063 | 0.6343 |
| KAD4_HUMAN | 3NDP (B)<sup>70</sup> | 2AR7 (A)<sup>71</sup> | 0.5371 | 0.5790 | 0.5913 | 0.6758 |
| KAD2_YEAST | 2AKY (A)<sup>72</sup> | 1DVR (B)<sup>73</sup> | 0.6795 | 0.7779 | 0.6795 | 0.7779 |

Overall, FingerprintContacts reliably predicts the open conformation for all the eight proteins, with template modeling (TM) scores of 0.6-0.8 in reference to the open crystal structure (Table 4). As a control, we refolded the structures by removing contacts unique to the closed state (Table S4, Figure S5). In all the eight test cases, FingerprintContacts outperforms the baseline results, with TM_open increase by 0.09 on average (Table S4, Figure S5). Moreover, FingerprintContacts successfully identified the open state for LIVK_ECOLI, RBSB_ECOLI, and KAD2_YEAST, which were not achieved by excluding all the contacts unique to the closed state (Figure S5). This suggests that FingerprintContacts may detect additional contacts that are crucial for the structural rearrangements from closed to open state. To further probe the predicted new structure, we superimposed the predicted structure onto the two available crystal structures. We find significant structural similarity between the predicted new structures and the open conformation (Figure 4, Figure S1). Specifically, we observe the dramatic domain opening in the predicted structure with respect to the experimental closed state while conserving the same topological features.

**Figure 4:**
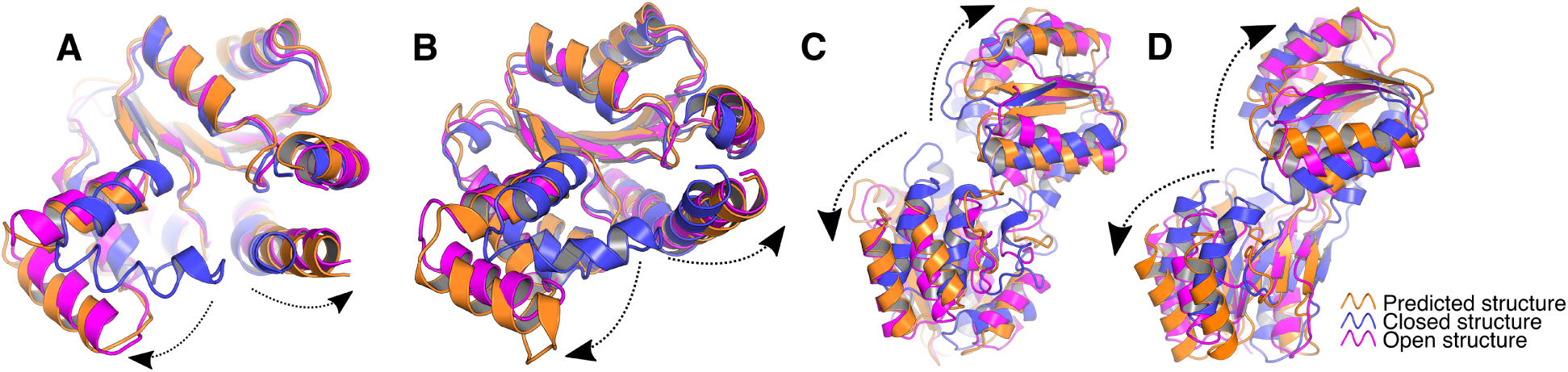
Structural superposition of predicted new structure against both conformers of a protein. (A) KAD_AQUAE, (B) KAD_ECOLI, (C) LIVK_ECOLI, and RBSB_ECOLI. Predicted open structure, closed, and open structures are represented in orange, blue, and magenta. The black lines represent the conformational motions.

Taken together, these results demonstrate that FingerprintContacts provides a powerful and automatic way to predict alternative conformations (open and closed state) using residue-residue contact information alone.

### FingerprintContacts identifies small communities of dynamically important residues

As FingerprintContacts outperforms the baseline results achieved by simply removing the conformation specific contacts, we wondered what type of contacts were removed by the algorithm. Many proteins undergo large-scale structural rearrangements through concerted motions of residues. As a result, we reasoned that the removed contacts could potentially correlate with the conformational fluctuations.^77^ To assess this, we investigated the relationship between removed contacts and moving residues, characterized by the changes of *C*_*β*_ distances (Δ*dist*_*C*_*β*_) in the alternative protein conformations (Figure 5). We observe a small number of clusters of residue-residue contacts, which is consistent with our hypothesis that a small set of contacts can be removed to identify the alternative conformation. Consistent with our expectation, our results suggest that the removed contacts are likely to represent residue pairs that experience large fluctuations (large Δ*dist*_*C*_*β*_). Another interesting observation is that there is a small fraction of removed contacts that are not true contacts in either conformations, termed as false positives previously (Figure 5A, C, D). Remarkably, we find that these removed false positives are actually spatially clustered with conformation specific contacts in the open state and have large Δ*dist*_*C*_*β*_ values. These “false positives” could exist in other intermediate conformational states and be crucial for the conformational transitions. ^78^ This could potentially explain why removing conformation specific contacts in the closed state alone was not sufficient to predict the open conformation (Figure S5).

**Figure 5:**
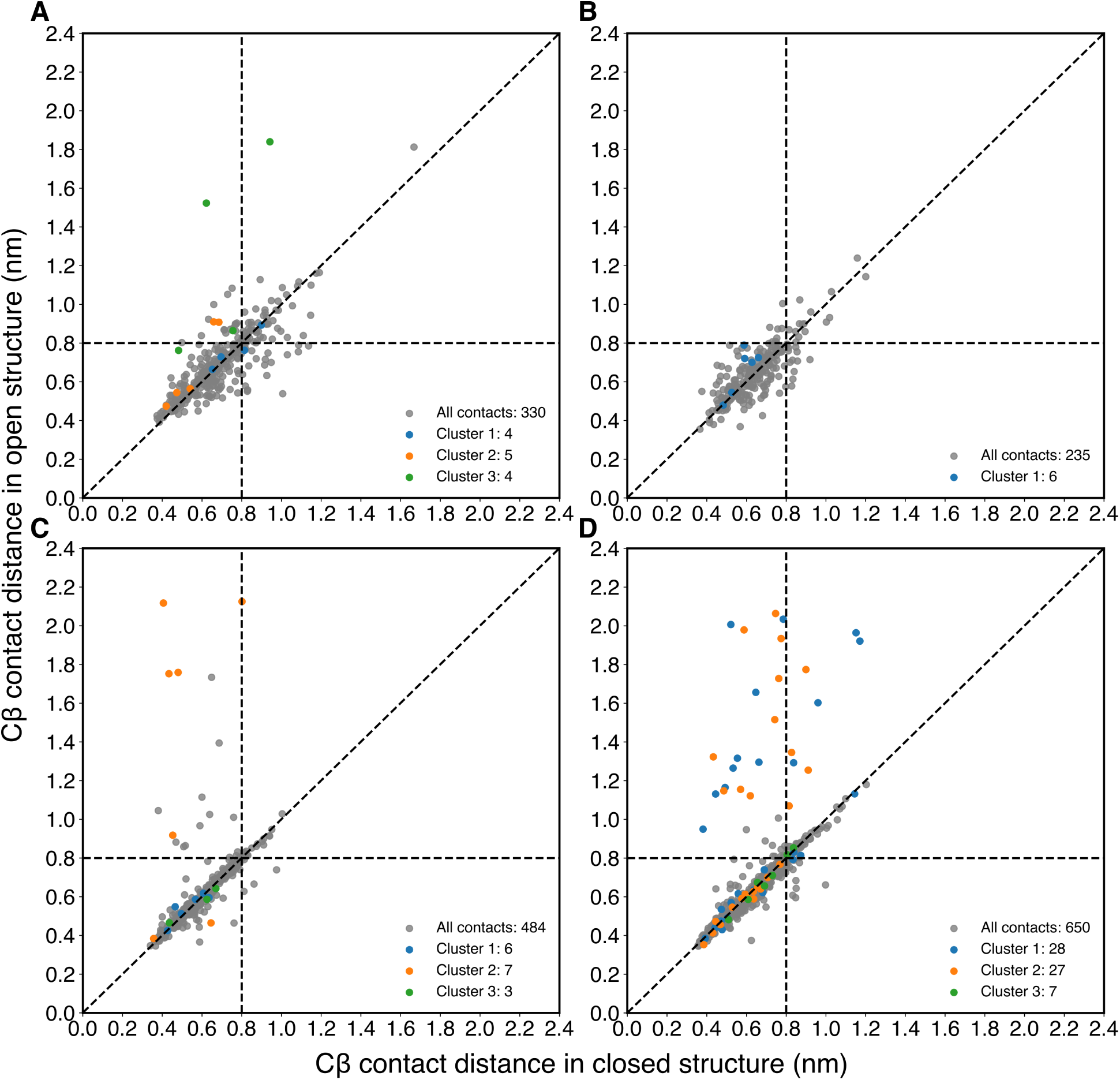
C_*β*_ distance distribution of removed contacts in both conformers. (A) KAD_AQUAE, (B) KAD_ECOLI, (C) LIVK_ECOLI, and (D) RBSB_ECOLI. The horizontal and vertical dashed lines represent the cutoff (8 Å), which is used to define whether a contact is true or not. The diagonal dashed line represents the case when *C*_*β*_ distances are the same in both open and closed structures. All the *default contacts* are shown in gray. The removed clusters of residue-residue contacts are marked in blue, orange, and green.

To better illustrate the correlation of the removed contacts and dynamically flexible regions, we take KAD_AQUAE as an example. We mapped the removed clusters of residueresidue contacts on the protein structures (Figure 6 and Figure S6-8). We observe that contacts removed in FingerprintContacts are preferentially located in the functionally dynamic regions. This insight could be of great use for extracting protein dynamics information from protein sequence. Previous studies have shown that coevolving residue pairs provide information about the spatial contacts within the 3D protein structure, conformational diversity, ligand binding, and protein oligomerization.^79^ However, it has been challenging to distinguish different signals of residue-residue contacts. Our results highlight the value of FingerprintContacts for identifying dynamically relevant residue-residue contacts. These discovered contacts could also aid in the *a priori* prediction of collective variables^80^ for molecular dynamics (MD) simulations, a powerful tool to characterize the conformational dynamics of proteins. Currently, the capabilities of MD simulations are severely constrained by two major problems: (1) sufficient sampling and (2) interpretation of high dimensional MD data. The detected residue-residue contacts could help in both cases by serving as collective variables to enhance the sampling or to represent the high dimensional MD data for human insights.

**Figure 6:**
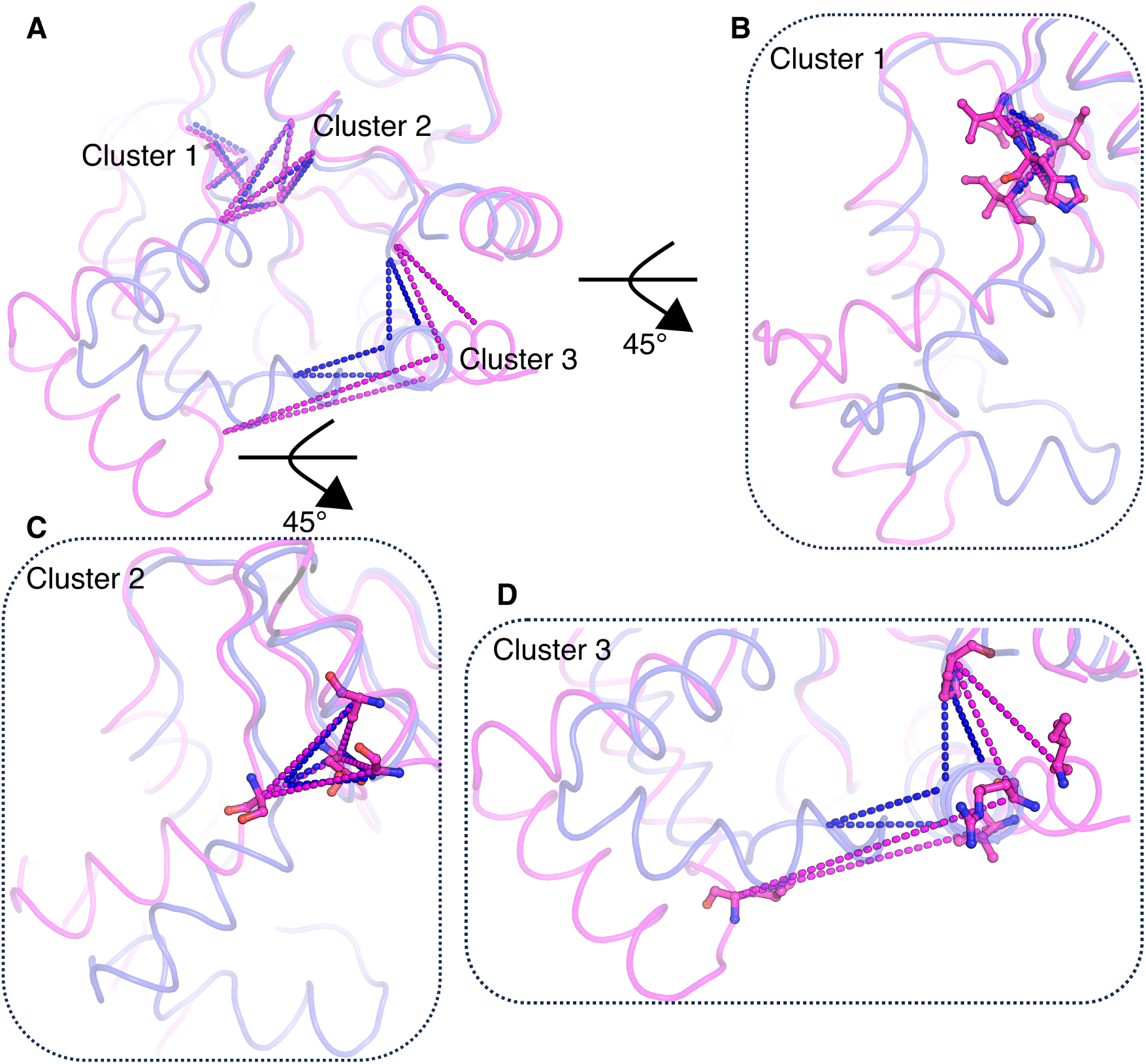
Illustration of detailed analysis for KAD_AQUAE. (A) Visualization of the three detected clusters in both closed (PDB: 3SR0_B^60^) and open (PDB: 2RH5_A^61^) crystal structures. (B-D) Zoom in view of Cluster 1, 2, and 3. The open structure is superpositioned onto the closed structure.

## Conclusions

We have introduced FingerprintContacts, a simple and efficient procedure to predict alternative protein conformations from residue-residue contacts alone. This algorithm was inspired by our previous work that few residue-residue contacts suffice to capture the complex conformational dynamics associated with protein folding and conformational changes. ^81–83^ We have validated our approach on eight ligand binding proteins with varying conformational movements. These proteins adopt two structurally distinct conformations: open and closed state. Analysis of the *default structure*, structure maximally satisfying the residue-residue contacts, and the *default contacts*, the corresponding subset of residue-residue contacts, provides two valuable insights: (1) the *default structure* strongly resembles the closed conformer despite the existence of similar number of conformation specific contacts; and (2) conformation specific contacts are only a small fraction of the *default contacts* (around 5 %) and rank lower among the *default contacts*, which results in the similar contact satisfaction score between the open and closed states. We leverage these observations to (1) predict the *default structure* as the closed conformer, (2) explore a small set of residue-residue contacts that perturb the *default structure* while satisfying similar amount of the *default contacts*, and (3) explore the alternative open conformation by removing these contacts from the structure prediction constraints.

To test FingerprintContacts, we compared the predicted structures with experimental alternative conformations and compared its performance to the previously reported approach, which refolds the structure by removing residue-residue contacts unique to the closed conformation. In all the eight cases, FingerprintContacts outperforms the baseline results, with an increase in TMscore by 0.09 on average. It is worth mentioning that no structural information is required for FingerprintContacts, whereas the baseline approach identifies the conformation specific contacts by comparing alternative conformations. The superposition of the predicted open structure shows significant similarity with the experimental open conformation. Moreover, the residue-residue contacts removed by FingerprintContacts shows clear correlation with dynamically fluctuating residues. Compared with current methods, FingerprintContacts provides two clear advantages: (1) widest scope, being applicable to proteins with only sequence information available, and (2) computational efficiency, being able to avoid the long-timescale simulations.

Because proteins perform diverse functions by adopting different conformations,^81,82^ we expect FingerprintContacts to serve as a powerful and cheap first step in the analysis of protein functional mechanisms. Beyond the prediction of alternative conformations, FingerprintContacts has provided important insights about the dynamically relevant residueresidue contacts (termed fingerprint contacts), offering many practical implications. With the rapid growth in protein sequence databases driven by next-generation sequencing, we expect FingerprintContacts will be widely applied to obtain a first glimpse of protein conformational diversity.

## Supplementary Information

Sample input file for FingerprintContacts.

Table S1: Parameters used in FingerprintContacts.

Table S2: Additional information about the *default structure*.

Table S3: Additional information about FingerprintContacts results.

Table S4: Comparison of baseline structure against *default structure* and both conformers of a protein.

Figure S1: Structural superposition of default and predicted structure against both conformers of AROA_ECOLI, ARGT_SALTY, KAD4_HUMAN, and KAD2_YEAST.

Figure S2: Distribution of conformation specific contacts in the alternative conformers of AROA_ECOLI, ARGT_SALTY, KAD4_HUMAN, and KAD2_YEAST.

Figure S3: Comparison between weighted contact satisfaction scores (*Q*_*s*_) in alternative conformers of the same protein.

Figure S4: Comparison between the predicted open structure and the *default structure*.

Figure S5: Structural similarity of predicted and baseline structures against the alternative conformations.

Figure S6: Illustration of fingerprint contacts for KAD_ECOLI.

Figure S7: Illustration of fingerprint contacts for LIVK_ECOLI.

Figure S8: Illustration of fingerprint contacts for RBSB_ECOLI.

## Acknowledgement

We thank Jiming Chen and Matthew Chan from Shukla research group for the critical reading of this manuscript and for creating the cover image for this article respectively. We also thank the Blue Waters sustained-petascale computing project, which is supported by the National Science Foundation (Awards OCI-0725070 and ACI-1238993) and the state of Illinois. D.S. acknowledges support from the New Innovator Award from the Foundation for Food and Agriculture Research and NSF Early Career Award (MCB 1845606). J.F. was partially supported by Chia-chen Chu Fellowship and Harry G. Drickamer Graduate Research Fellowship from University of Illinois at Urbana-Champaign.

## References

(1) Dror, R. O.; Arlow, D. H.; Maragakis, P.; Mildorf, T. J.; Pan, A. C.; Xu, H.; Borhani, D. W.; Shaw, D. E. Activation Mechanism of the β2-adrenergic Receptor. Proc. Natl. Acad. Sci. USA 2011, 108, 18684–18689.

(2) Dror, R. O.; Pan, A. C.; Arlow, D. H.; Borhani, D. W.; Maragakis, P.; Shan, Y.; Xu, H.; Shaw, D. E. Pathway and Mechanism of Drug Binding to G-protein-coupled Receptors. Proc. Natl. Acad. Sci. USA 2011, 108, 13118–13123.

(3) Shan, Y.; Kim, E. T.; Eastwood, M. P.; Dror, R. O.; Seeliger, M. A.; Shaw, D. E. How Does a Drug Molecule Find its Target Binding Site? J. Am. Chem. Soc. 2011, 133, 9181–9183.

(4) Lawrenz, M.; Shukla, D.; Pande, V. S. Cloud computing approaches for prediction of ligand binding poses and pathways. Sci. Rep. 2015, 5.

(5) Moffett, A. S.; Bender, K. W.; Huber, S. C.; Shukla, D. Allosteric Control of a Plant Receptor Kinase through S-glutathionylation. Biophys. J. 2017, 113, 2354–2363.

(6) Moffett, A. S.; Shukla, D. Using Molecular Simulation to Explore the Nanoscale Dynamics of the Plant Kinome. Biochem. J. 2018, 475, 905–921.

(7) Shukla, D.; Meng, Y.; Roux, B.; Pande, V. S. Activation Pathway of Src Kinase Reveals Intermediate States as Targets for Drug Design. Nat. Commun. 2014, 5, 3397.

(8) Selvam, B.; Mittal, S.; Shukla, D. Free Energy Landscape of the Complete Transport Cycle in a Key Bacterial Transporter. ACS Cent. Sci. 2018, 4, 1146–1154.

(9) Selvam, B.; Yu, Y.-C.; Chen, L.-Q.; Shukla, D. Molecular Basis of the Glucose Transport Mechanism in Plants. ACS Cent. Sci. 2019,

(10) Cheng, K. J.; Selvam, B.; Chen, L.-Q.; Shukla, D. Distinct Substrate Transport Mechanism Identified in Homologous Sugar Transporters. J. Phys. Chem. B 2019, 123, 8411–8418.

(11) Rigden, D. J.; Rigden, D. J. From Protein Structure to Function with Bioinformatics; Springer, 2009.

(12) Henzler-Wildman, K.; Kern, D. Dynamic Personalities of Proteins. Nature 2007, 450, 964.

(13) Mittal, S.; Shukla, D. Recruiting Machine Learning Methods for Molecular Simulations of Proteins. Mol. Simul. 2018, 44, 891–904.

(14) Schuler, B. Single-molecule FRET of Protein Structure and Dynamics - A Primer. J. Nanobiotechnol. 2013, 11, S2.

(15) Jeschke, G. DEER Distance Measurements on Proteins. Annu. Rev. Phys. Chem. 2012, 63, 419–446.

(16) Dror, R. O.; Dirks, R. M.; Grossman, J.; Xu, H.; Shaw, D. E. Biomolecular Simulation: A Computational Microscope for Molecular Biology. Annu. Rev. Biophys. 2012, 41, 429–452.

(17) UniProt: The Universal Protein Knowledgebase. Nucleic Acids Res. 2016, 45, D158–D169.

(18) Berman, H. M.; Westbrook, J.; Feng, Z.; Gilliland, G.; Bhat, T.; Weissig, H.; Shindyalov, I.; Bourne, P. The Protein Data Bank. Nucleic Acids Res 2000, 28, 235–242.

(19) Feng, J.; Chen, J.; Selvam, B.; Shukla, D. Computational Microscopy: Revealing Molecular Mechanisms in Plants using Mlecular Dynamics Simulations. Plant Cell 2019, 31, tpc.119.tt1219.

(20) Morcos, F.; Pagnani, A.; Lunt, B.; Bertolino, A.; Marks, D. S.; Sander, C.; Zecchina, R.; Onuchic, J. N.; Hwa, T.; Weigt, M. Direct-Coupling Analysis of Residue Coevolution Captures Native Contacts across many Protein Families. Proc. Natl. Acad. Sci. USA 2011, 108, E1293–E1301.

(21) Marks, D. S.; Colwell, L. J.; Sheridan, R.; Hopf, T. A.; Pagnani, A.; Zecchina, R.; Sander, C. Protein 3D Structure Computed from Evolutionary Sequence Variation. PLoS One 2011, 6, e28766.

(22) Marks, D. S.; Hopf, T. A.; Sander, C. Protein Structure Prediction from Sequence Variation. Nat. Biotechnol. 2012, 30, 1072–1080.

(23) Hopf, T. A.; Colwell, L. J.; Sheridan, R.; Rost, B.; Sander, C.; Marks, D. S. Three-Dimensional Structures of Membrane Proteins From Genomic Sequencing. Cell 2012, 149, 1607–1621.

(24) Hopf, T. A.; Schärfe, C. P.; Rodrigues, J. P.; Green, A. G.; Kohlbacher, O.; Sander, C.; Bonvin, A. M.; Marks, D. S. Sequence Co-evolution Gives 3D Contacts and Structures of Protein Complexes. Elife 2014, 3, e03430.

(25) Kryshtafovych, A.; Schwede, T.; Topf, M.; Fidelis, K.; Moult, J. Critical Assessment of Methods of Protein Structure Prediction (CASP) - Round XIII. Proteins 2019,

(26) Simkovic, F.; Ovchinnikov, S.; Baker, D.; Rigden, D. J. Applications of Contact Predictions to Structural Biology. IUCrJ 2017, 4, 291–300.

(27) Jones, D. T.; Buchan, D. W.; Cozzetto, D.; Pontil, M. PSICOV: Precise Structural Contact Prediction using Sparse Inverse Covariance Estimation on Large Multiple Sequence Alignments. Bioinformatics 2011, 28, 184–190.

(28) Seemayer, S.; Gruber, M.; Söding, J. CCMpred - Fast and Precise Prediction of Protein Residue - Residue Contacts from Correlated Mutations. Bioinformatics 2014, 30, 3128–3130.

(29) Ovchinnikov, S.; Kinch, L.; Park, H.; Liao, Y.; Pei, J.; Kim, D. E.; Kamisetty, H.; Grishin, N. V.; Baker, D. Large-scale Determination of Previously Unsolved Protein Structures using Evolutionary Information. Elife 2015, 4, e09248.

(30) Lensink, M. F.; Brysbaert, G.; Nadzirin, N.; Velankar, S.; Chaleil, R. A.; Gerguri, T.; Bates, P. A.; Laine, E.; Carbone, A.; Grudinin, S. et al. Blind Prediction of Homo- and Hetero-protein Complexes: The CASP13-CAPRI Experiment. Proteins 2019,

(31) Weinreb, C.; Riesselman, A. J.; Ingraham, J. B.; Gross, T.; Sander, C.; Marks, D. S. 3D RNA and Functional Interactions from Evolutionary Couplings. Cell 2016, 165, 963–975.

(32) Hopf, T. A.; Ingraham, J. B.; Poelwijk, F. J.; Schärfe, C. P.; Springer, M.; Sander, C.; Marks, D. S. Mutation Effects Predicted from Sequence Co-variation. Nat. Biotechnol. 2017, 35, 128.

(33) Ovchinnikov, S.; Park, H.; Varghese, N.; Huang, P.-S.; Pavlopoulos, G. A.; Kim, D. E.; Kamisetty, H.; Kyrpides, N. C.; Baker, D. Protein Structure Determination using Metagenome Sequence Data. Science 2017, 355, 294–298.

(34) Wu, S.; Zhang, Y. A Comprehensive Assessment of Sequence-based and Template-based Methods for Protein Contact Prediction. Bioinformatics 2008, 24, 924–931.

(35) Cheng, J.; Randall, A. Z.; Sweredoski, M. J.; Baldi, P. SCRATCH: A Protein Structure and Structural Feature Prediction Server. Nucleic Acids Res. 2005, 33, W72–W76.

(36) Skwark, M. J.; Raimondi, D.; Michel, M.; Elofsson, A. Improved Contact Predictions using the Recognition of Protein Like Contact Patterns. PLoS Comput. Biol. 2014, 10, e1003889.

(37) Jones, D. T.; Singh, T.; Kosciolek, T.; Tetchner, S. MetaPSICOV: Combining Coevolution Methods for Accurate Prediction of Contacts and Long Range Hydrogen Bonding in Proteins. Bioinformatics 2014, 31, 999–1006.

(38) Wang, Z.; Xu, J. Predicting Protein Contact Map using Evolutionary and Physical Constraints by Integer Programming. Bioinformatics 2013, 29, i266–i273.

(39) Ma, J.; Wang, S.; Wang, Z.; Xu, J. Protein Contact Prediction by Integrating Joint Evolutionary Coupling Analysis and Supervised Learning. Bioinformatics 2015, 31, 3506–3513.

(40) Wang, S.; Sun, S.; Li, Z.; Zhang, R.; Xu, J. Accurate De Novo Prediction of Protein Contact Map by Ultra-deep Learning Model. PLoS Comput. Biol. 2017, 13, e1005324.

(41) de Oliveira, S. H. P.; Shi, J.; Deane, C. M. Comparing Co-evolution Methods and their Application to Template-free Protein Structure Prediction. Bioinformatics 2017, 33, 373–381.

(42) Wuyun, Q.; Zheng, W.; Peng, Z.; Yang, J. A Large-scale Comparative Assessment of Methods for Residue-Residue Contact Prediction. Brief. Bioinform. 2016, 19, 219–230.

(43) Kuhlman, B.; Bradley, P. Advances in Protein Structure Prediction and Design. Nature Reviews Molecular Cell Biology 2019, 1–17.

(44) Zea, D. J.; Monzon, A. M.; Parisi, G.; Marino-Buslje, C. How is Structural Divergence Related to Evolutionary Information? Mol. Phylogenetics Evol. 2018, 127, 859–866.

(45) Morcos, F.; Jana, B.; Hwa, T.; Onuchic, J. N. Coevolutionary Signals across Protein Lineages Help Capture Multiple Protein Conformations. Proc. Natl. Acad. Sci. USA 2013, 110, 20533–20538.

(46) Sutto, L.; Marsili, S.; Valencia, A.; Gervasio, F. L. From Residue Coevolution to Protein Conformational Ensembles and Functional Dynamics. Proc. Natl. Acad. Sci. USA 2015, 112, 13567–13572.

(47) Sfriso, P.; Duran-Frigola, M.; Mosca, R.; Emperador, A.; Aloy, P.; Orozco, M. Residues Coevolution Guides the Systematic Identification of Alternative Functional Conformations in Proteins. Structure 2016, 24, 116–126.

(48) Shamsi, Z.; Moffett, A. S.; Shukla, D. Enhanced Unbiased Sampling of Protein Dynamics using Evolutionary Coupling Information. Sci. Rep. 2017, 7, 12700.

(49) Feng, J.; Shukla, D. Characterizing Conformational Dynamics of Proteins using Evolutionary Couplings. J. Phys. Chem. B 2018, 122, 1017–1025.

(50) Shamsi, Z.; Cheng, K. J.; Shukla, D. Reinforcement Learning Based Adaptive Sampling: REAPing Rewards by Exploring Protein Conformational Landscapes. J. Phys. Chem. B 2018, 122, 8386–8395.

(51) Granata, D.; Ponzoni, L.; Micheletti, C.; Carnevale, V. Patterns of Coevolving Amino Acids Unveil Structural and Dynamical Domains. Proc. Natl. Acad. Sci. USA 2017, 114, E10612–E10621.

(52) Halabi, N.; Rivoire, O.; Leibler, S.; Ranganathan, R. Protein Sectors: Evolutionary Units of Three-dimensional Structure. Cell 2009, 138, 774–786.

(53) Rokach, L.; Maimon, O. Data Mining and Knowledge Discovery Handbook; Springer, 2005; pp 321–352.

(54) Oliphant, T. E. A Guide to NumPy; Trelgol Publishing USA, 2006; Vol. 1.

(55) McKinney, W. Data Structures for Statistical Computing in Python. Proc. 9th Python Sci. Conf. 2010, 445, 51–56.

(56) Pedregosa, F.; Varoquaux, G.; Gramfort, A.; Michel, V.; Thirion, B.; Grisel, O.; Blondel, M.; Prettenhofer, P.; Weiss, R.; Dubourg, V. et al. Scikit-learn: Machine Learning in Python. JMLR 2011, 12, 2825–2830.

(57) Adhikari, B.; Cheng, J. CONFOLD2: Improved Contact-driven Ab Initio Protein Structure Modeling. BMC Bioinform. 2018, 19, 22.

(58) Xu, J.; Zhang, Y. How Significant is a Protein Structure Similarity with TM-score = 0.5? Bioinformatics 2010, 26, 889–895.

(59) Kreyszig, E. Advanced Engineering Mathematics. Fourth Edi. 1979.

(60) Kerns, S. J.; Agafonov, R. V.; Cho, Y.-J.; Pontiggia, F.; Otten, R.; Pachov, D. V.; Kutter, S.; Phung, L. A.; Murphy, P. N.; Thai, V. et al. The Energy Landscape of Adenylate Kinase during Catalysis. Nat. Struct. Mol. Biol. 2015, 22, 124.

(61) Henzler-Wildman, K. A.; Thai, V.; Lei, M.; Ott, M.; Wolf-Watz, M.; Fenn, T.; Pozharski, E.; Wilson, M. A.; Petsko, G. A.; Karplus, M. et al. Intrinsic Motions along an Anzymatic Reaction Trajectory. Nature 2007, 450, 838.

(62) Müller, C. W.; Schulz, G. E. Structure of the Complex between Adenylate Kinase from Escherichia Coli and the Inhibitor Ap5A Refined at 1.9 Å Resolution: A Model for a Catalytic Transition State. J. Mol. Biol. 1992, 224, 159–177.

(63) Müller, C.; Schlauderer, G.; Reinstein, J.; Schulz, G. E. Adenylate Kinase Motions during Catalysis: An Energetic Counterweight Balancing Substrate Binding. Struc. 1996, 4, 147–156.

(64) Magnusson, U.; Salopek-Sondi, B.; Luck, L. A.; Mowbray, S. L. X-ray Structures of the Leucine-binding Protein Illustrate Conformational Changes and the Basis of Ligand Specificity. J. Biol. Chem. 2004, 279, 8747–8752.

(65) Björkman, A.; Binnie, R. A.; Zhang, H.; Cole, L. B.; Hermodson, M. A.; Mowbray, S. L. Probing Protein-Protein Interactions. The Ribose-Binding Protein in Bacterial Transport and Chemotaxis. J. Biol. Chem. 1994, 269, 30206–30211.

(66) Björkman, A. J.; Mowbray, S. L. Multiple Open Forms of Ribose-Binding Protein Trace the Path of its Conformational Change. J. Mol. Biol. 1998, 279, 651–664.

(67) Priestman, M. A.; Healy, M. L.; Funke, T.; Becker, A.; Schönbrunn, E. Molecular Basis for the Glyphosate-insensitivity of the Reaction of 5-enolpyruvylshikimate 3-phosphate Synthase with Shikimate. FEBS Lett. 2005, 579, 5773–5780.

(68) Stallings, W. C.; Abdel-Meguid, S. S.; Lim, L. W.; Shieh, H. S.; Dayringer, H. E.; Leimgruber, N. K.; Stegeman, R. A.; Anderson, K. S.; Sikorski, J. A.; Padgette, S. R. et al. Structure and Topological Symmetry of the Glyphosate Target 5-enolpyruvylshikimate-3-phosphate Synthase: A Distinctive Protein Fold. Proc. Natl. Acad. Sci. USA 1991, 88, 5046–5050.

(69) Oh, B.-H.; Pandit, J.; Kang, C.-H.; Nikaido, K.; Gokcen, S.; Ames, G.; Kim, S.-H. Three-dimensional Structures of the Periplasmic Lysine/arginine/ornithine-binding Protein with and without a Ligand. J. Biol. Chem. 1993, 268, 11348–11355.

(70) Liu, R.; Xu, H.; Wei, Z.; Wang, Y.; Lin, Y.; Gong, W. Crystal Structure of Human Adenylate Kinase 4 (L171P) Suggests the Role of Hinge Region in Protein Domain Motion. Biochem. Biophys. Res. Commun. 2009, 379, 92–97.

(71) Filippakopoulos, P.; Turnbull, A.; Fedorov, O.; Weigelt, J.; Bunkoczi, G.; Ugochukwu, E.; Debreczeni, J.; Niesen, F.; von Delft, F.; Edwards, A. et al. Crystal Structure of Human Adenylate Kinase 4, AK4. 2005; https://doi.org/10.2210/pdb2ar7/pdb.

(72) Abele, U.; Schulz, G. High-resolution Structures of Adenylate Kinase from Yeast Ligated with Inhibitor Ap5A, Showing the Pathway of Phosphoryl Transfer. Protein Sci. 1995, 4, 1262–1271.

(73) Schlauderer, G.; Proba, K.; Schulz, G. Structure of a Mutant Adenylate Kinase Ligated with an ATP-analogue Showing Domain Closure Over ATP. J. Mol. Biol. 1996, 256, 223–227.

(74) Wei, G.; Xi, W.; Nussinov, R.; Ma, B. Protein Ensembles: How Does Nature Harness Thermodynamic Fluctuations for Life? The Diverse Functional Roles of Conformational Ensembles in the Cell. Chem. Rev. 2016, 116, 6516–6551.

(75) Uguzzoni, G.; Lovis, S. J.; Oteri, F.; Schug, A.; Szurmant, H.; Weigt, M. Large-scale Identification of Coevolution Signals across Homo-oligomeric Protein Interfaces by Direct Coupling Analysis. Proc. Natl. Acad. Sci. USA 2017, 114, E2662–E2671.

(76) Tinberg, C. E.; Khare, S. D.; Dou, J.; Doyle, L.; Nelson, J. W.; Schena, A.; Jankowski, W.; Kalodimos, C. G.; Johnsson, K.; Stoddard, B. L. et al. Computational Design of Ligand-binding Proteins with High Affinity and Selectivity. Nature 2013, 501, 212.

(77) Jeon, J.; Nam, H.-J.; Choi, Y. S.; Yang, J.-S.; Hwang, J.; Kim, S. Molecular Evolution of Protein Conformational Changes Revealed by a Network of Evolutionarily Coupled Residues. Mol. Biol. Evol. 2011, 28, 2675–2685.

(78) Anishchenko, I.; Ovchinnikov, S.; Kamisetty, H.; Baker, D. Origins of Coevolution between Residues Distant in Protein 3D Structures. Proc. Natl. Acad. Sci. USA 2017, 114, 9122–9127.

(79) Hopf, T. A.; Marks, D. S. From Protein Structure to Function with Bioinformatics; Springer, 2017; pp 37–58.

(80) McGibbon, R. T.; Husic, B. E.; Pande, V. S. Identification of Simple Reaction Coordinates from Complex Dynamics. J. Chem. Phys. 2017, 146, 044109.

(81) Lane, T. J.; Shukla, D.; Beauchamp, K. A.; Pande, V. S. To milliseconds and beyond: challenges in the simulation of protein folding. Curr. Opin. Struc. Bio. 2013, 23, 58–65.

(82) Shukla, D.; HernÃąndez, C. X.; Weber, J. K.; Pande, V. S. Markov State Models Provide Insights into Dynamic Modulation of Protein Function. Acc. Chem. Res. 2015, 48, 414–422.

(83) Kohlhoff, K.; Shukla, D.; Lawrenz, M.; Bowman, G.; Konerding, D. E.; Belov, D.; B., A. R.; Pande, V. S. Cloud-based Simulations on Google Exacycle Reveal Ligand Modulation of GPCR Activation Pathways. Nat. Chem. 2014, 6, 15–21.

