## Supplementary Information for "FingerprintContacts: Predicting Alternative Conformations of Proteins from Coevolution"

### Supporting information for: FingerprintContacts: Predicting Alternative Conformations of Proteins from Coevolution

### Sample input file for FingerprintContacts

A sample input file for FingerprintContacts is provided. All the parameters used in FingerprintContacts are defined explicitly in the input file. Parameter values can be adjusted if found appropriate.

### Software Directory
sourceCodePath="/home/FingerprintContacts"
confoldPath="/home/CONFOLD2-master/confold-v2.0"
### Output Path
output_path="/home/results/1ba2"
### Protein Information
contact_file="/home/results/1ba2/1ba2.rr"
secondary_structure_file="/home/results/1ba2/1ba2.ss"
### Clustering Parameters
n_clusters_range=[10, 100, 10]
bound=[0.005, 0.05, 0.1]
### Scoring Parameters
tm_thres=0.5
Q_s_thres=0.6
### Selecting Parameters
zscore_thres=1.5
reward_thres=0.5
### Comparison (Optional)
structure_A="/home/results/1ba2/2dri.pdb"
structure_B="/home/results/1ba2/1ba2.pdb"

**Table S1: Parameters used in FingerprintContacts**

| Uniprot Name | Clustering Parameters | | Scoring Parameters | | Selecting Parameters | |
|---|---|---|---|---|---|---|
| | n_clusters_range | bound | tm_thres | Q_s_thres | zscore_thres | reward_thres |
| KAD_AQUAE | [10, 100, 10] | [0.005, 0.05, 0.05] | 0.5 | 0.65 | 1.5 | 0.5 |
| KAD_ECOLI | [10, 100, 10] | [0.005, 0.05, 0.1] | 0.5 | 0.7 | 1.5 | 0.5 |
| LIVK_ECOLI | [10, 100, 10] | [0.005, 0.05, 0.05] | 0.5 | 0.7 | 1.5 | 0.5 |
| RBSB_ECOLI | [10, 100, 10] | [0.005, 0.05, 0.1] | 0.5 | 0.6 | 1.5 | 0.5 |
| AROA_ECOLI | [10, 100, 10] | [0.005, 0.05, 0.1] | 0.5 | 0.7 | 1.5 | 0.5 |
| ARGT_SALTY | [10, 100, 10] | [0.005, 0.05, 0.05] | 0.5 | 0.7 | 1.5 | 0.5 |
| KAD4_HUMAN | [10, 100, 10] | [0.005, 0.05, 0.1] | 0.5 | 0.6 | 1.5 | 0.5 |
| KAD2_YEAST | [10, 100, 10] | [0.005, 0.05, 0.1] | 0.5 | 0.6 | 1.5 | 0.45 |

**Table S2: Additional information about the *default structure***

| Uniprot Name | Q_s | RMSD_closed (Å) | RMSD_open (Å) |
|---|---|---|---|
| KAD_AQUAE | 0.7363 | 4.534 | 6.035 |
| KAD_ECOLI | 0.8691 | 3.663 | 6.995 |
| LIVK_ECOLI | 0.9146 | 2.125 | 7.257 |
| RBSB_ECOLI | 0.7901 | 2.660 | 7.039 |
| AROA_ECOLI | 0.8183 | 4.298 | 7.251 |
| ARGT_SALTY | 0.8993 | 4.183 | 5.948 |
| KAD4_HUMAN | 0.8566 | 7.412 | 9.350 |
| KAD2_YEAST | 0.7793 | 3.235 | 5.196 |

**Table S3: Additional information about FingerprintContacts results**

| Uniprot Name | Blind top | | | | Best | | | |
|---|---|---|---|---|---|---|---|---|
| | #_Contacts | Reward | Q_s | TM_ini | #_Contacts | Reward | Q_s | TM_ini |
| KAD_AQUAE | 11 | 0.6221 | 0.6674 | 0.6165 | 13 | 0.5370 | 0.6927 | 0.6506 |
| KAD_ECOLI | 3 | 0.8242 | 0.7311 | 0.5482 | 6 | 0.6179 | 0.7261 | 0.6181 |
| LIVK_ECOLI | 24 | 0.9790 | 0.7181 | 0.5053 | 16 | 0.6375 | 0.7056 | 0.6107 |
| RBSB_ECOLI | 63 | 0.8298 | 0.7394 | 0.5465 | 62 | 0.6461 | 0.7295 | 0.6075 |
| AROA_ECOLI | 13 | 0.9980 | 0.7150 | 0.5005 | 5 | 0.6407 | 0.7353 | 0.6095 |
| ARGT_SALTY | 11 | 0.9135 | 0.7097 | 0.5226 | 4 | 0.5337 | 0.7371 | 0.6520 |
| KAD4_HUMAN | 15 | 0.9992 | 0.6201 | 0.5002 | 20 | 0.7489 | 0.7092 | 0.5718 |
| KAD2_YEAST | 49 | 0.4791 | 0.6743 | 0.6761 | 49 | 0.4791 | 0.6743 | 0.6761 |

**Table S4: Comparison of baseline structure against *default structure* and both conformers of a protein**

| Uniprot Name | TM_ini | TM_closed | TM_open |
|---|---|---|---|
| KAD_AQUAE | 0.7252 | 0.7040 | 0.7875 |
| KAD_ECOLI | 0.5419 | 0.5641 | 0.6895 |
| LIVK_ECOLI | 0.7966 | 0.7729 | 0.6065 |
| RBSB_ECOLI | 0.8895 | 0.8674 | 0.6396 |
| AROA_ECOLI | 0.5241 | 0.5049 | 0.6167 |
| ARGT_SALTY | 0.5171 | 0.5092 | 0.5350 |
| KAD4_HUMAN | 0.5991 | 0.5747 | 0.6189 |
| KAD2_YEAST | 0.8283 | 0.7640 | 0.6720 |

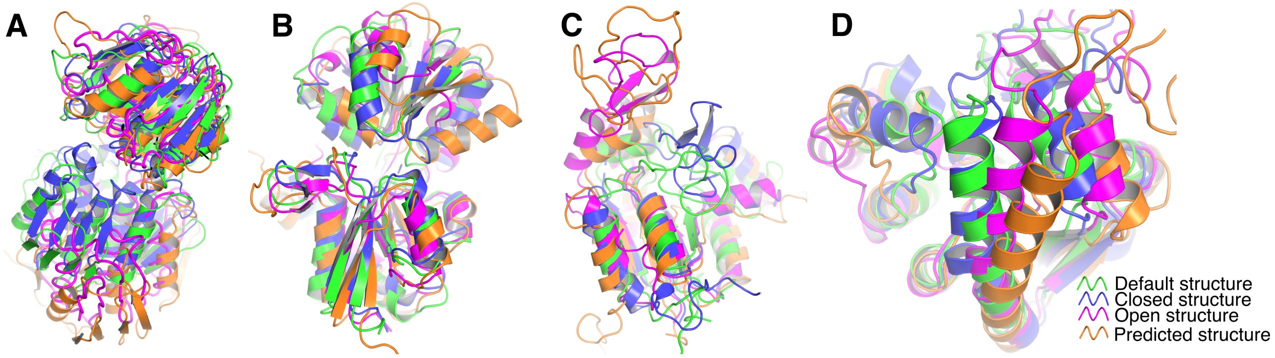


Figure S1: **Structural superposition of default and predicted structure against both conformers of a protein.** (A) AROA_ECOLI, (B) ARGT_SALTY, (C) KAD4_HUMAN, and (D) KAD2_YEAST. Default structure, closed, open, and predicted structures are represented in green, blue, magenta, and orange.

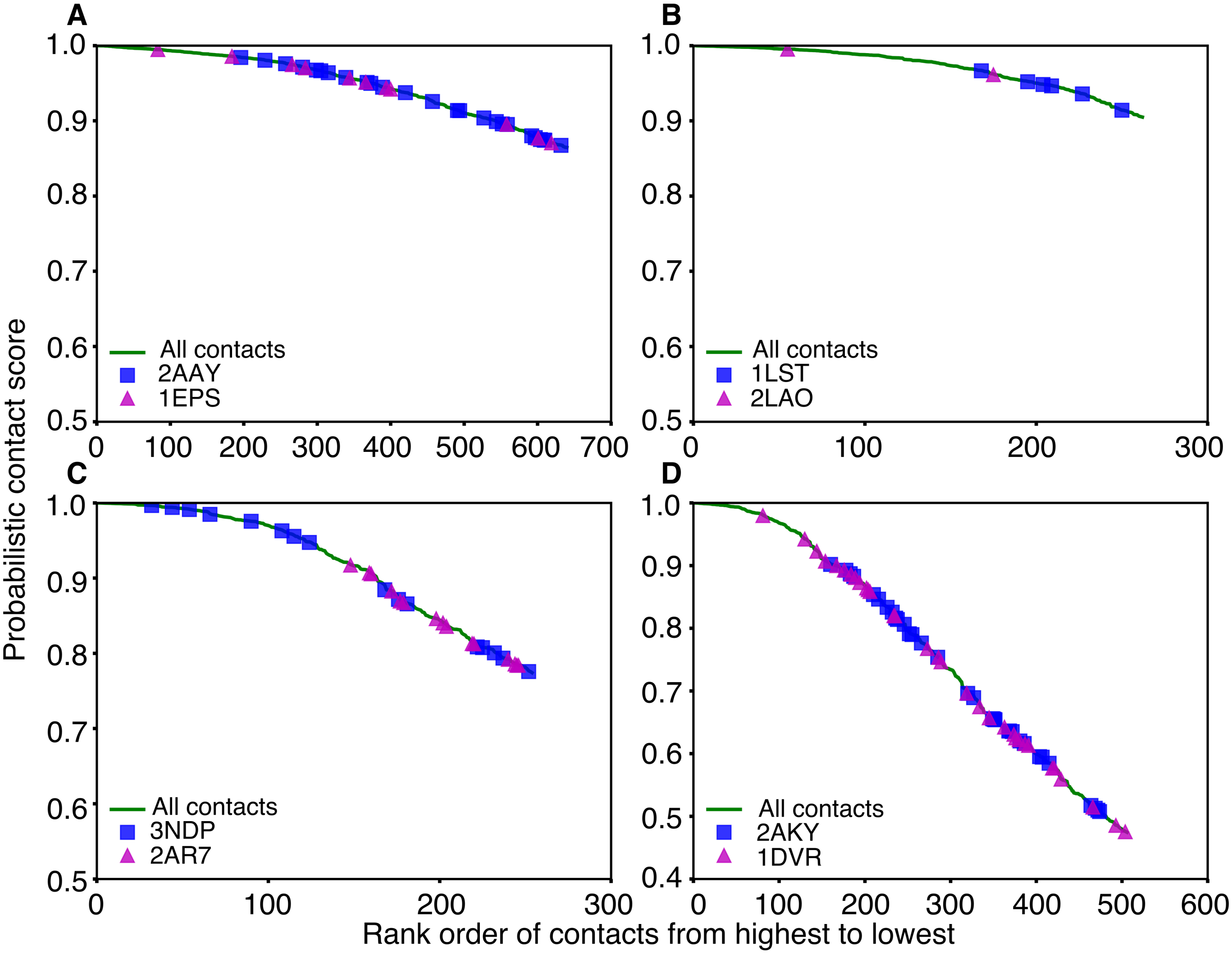


Figure S2: **Distribution of conformation specific contacts in the alternative conformers of a protein.** (A) AROA_ECOLI, (B) ARGT_SALTY, (C) KAD4_HUMAN, and (D) KAD2_YEAST. All the *default contacts*, residue-residue contacts unique to closed and open structures are shown in green, blue, and magenta, respectively.

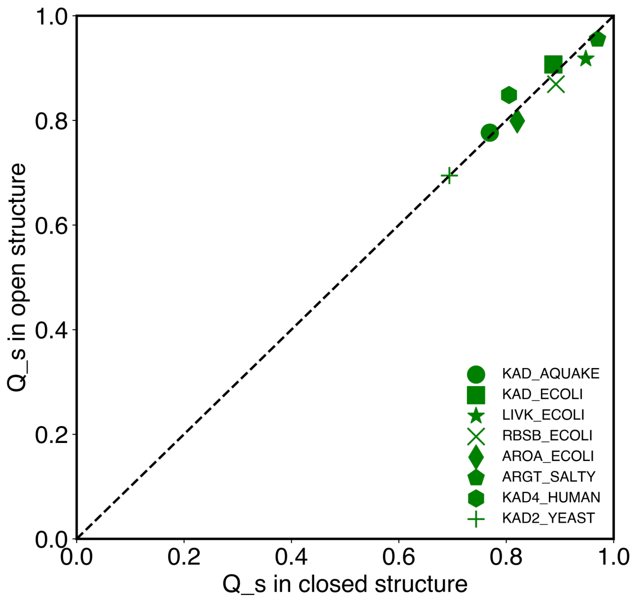


Figure S3: **Comparison between weighted contact satisfaction scores ($Q_s$) in alternative conformers of the same protein.** The dashed line indicates the case when $Q_s$ is the same in both structures.

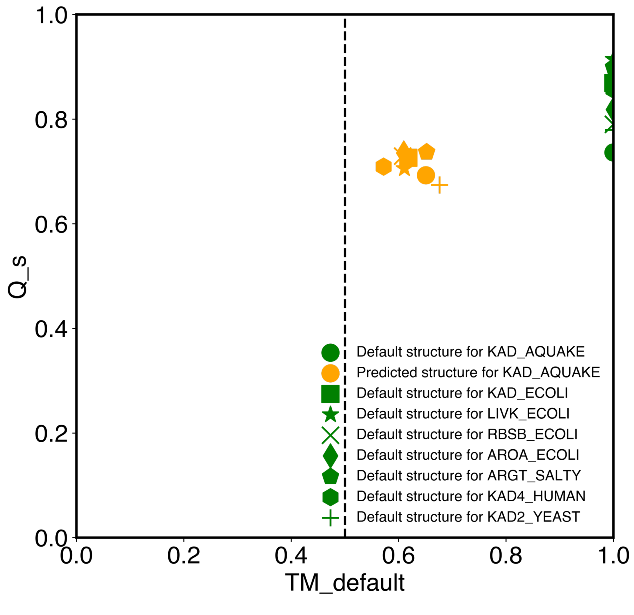


Figure S4: **Comparison between the predicted open structure and the *default* structure.** $Q_s$ represents the weighted contact satisfaction score and TM_default shows the TMscore between the two structures. The vertical dashed line indicates the chosen cutoff ($tm\_thres = 0.5$).

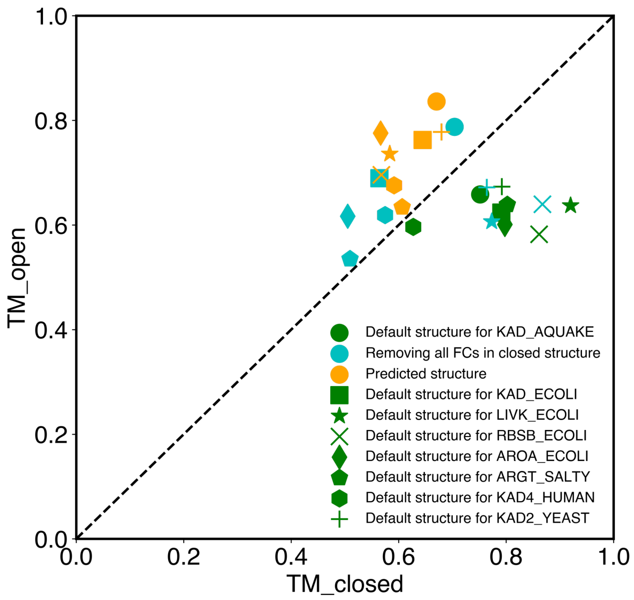


Figure S5: **Structural similarity of predicted and baseline structures against the alternative conformations.** TM_closed and TM_open represent the TMscore between the given structure and the closed and open conformations, respectively. The dashed line indicates the case when the structure is equally distant to the two conformations.

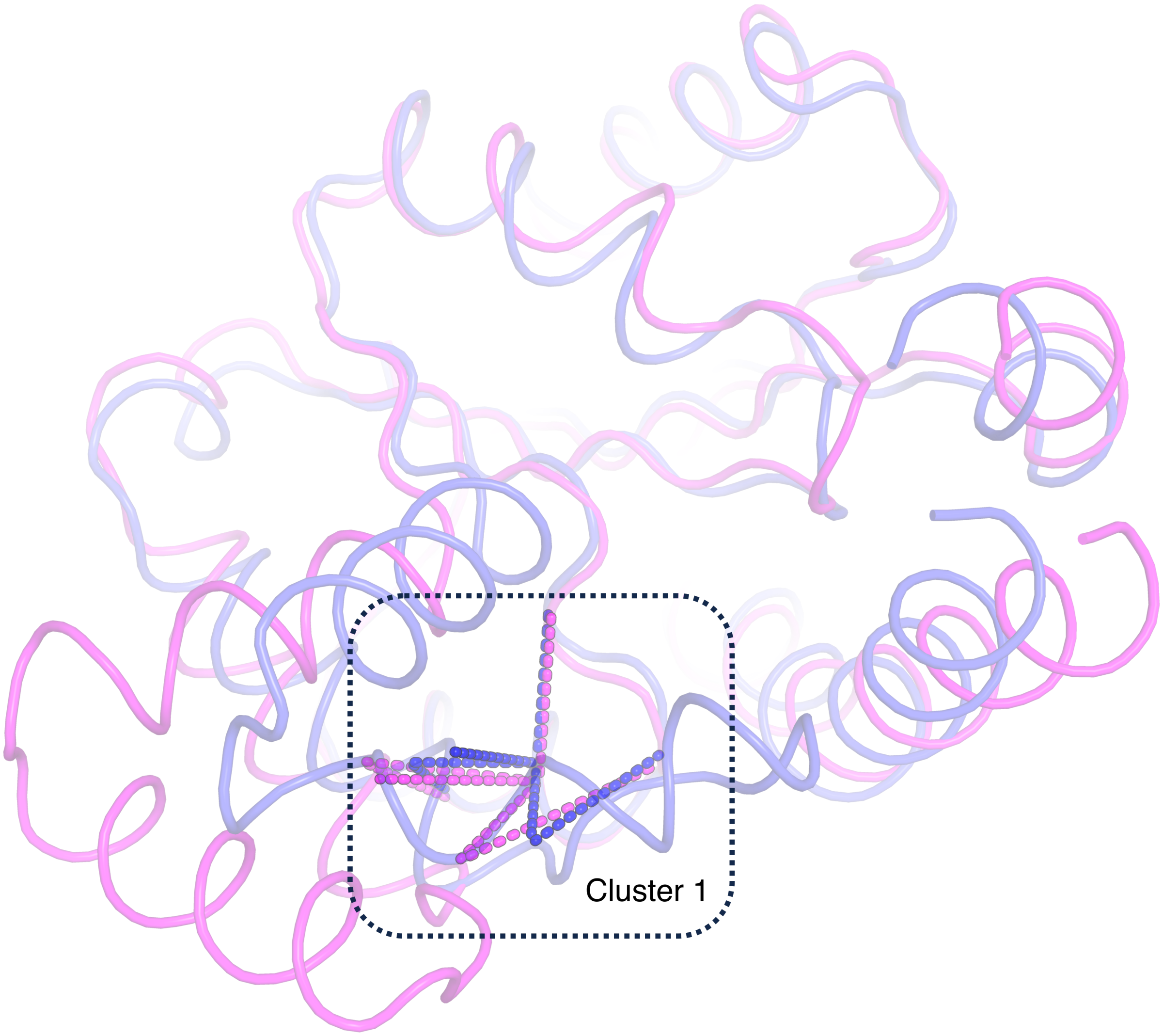


Figure S6: **Illustration of fingerprint contacts for KAD_ECOLI.** Visualization of the detected cluster in both closed (PDB: 1AKE_A [S1]) and open (PDB: 4AKE_A [S2]) crystal structures. The open structure is superimposed onto the closed structure.

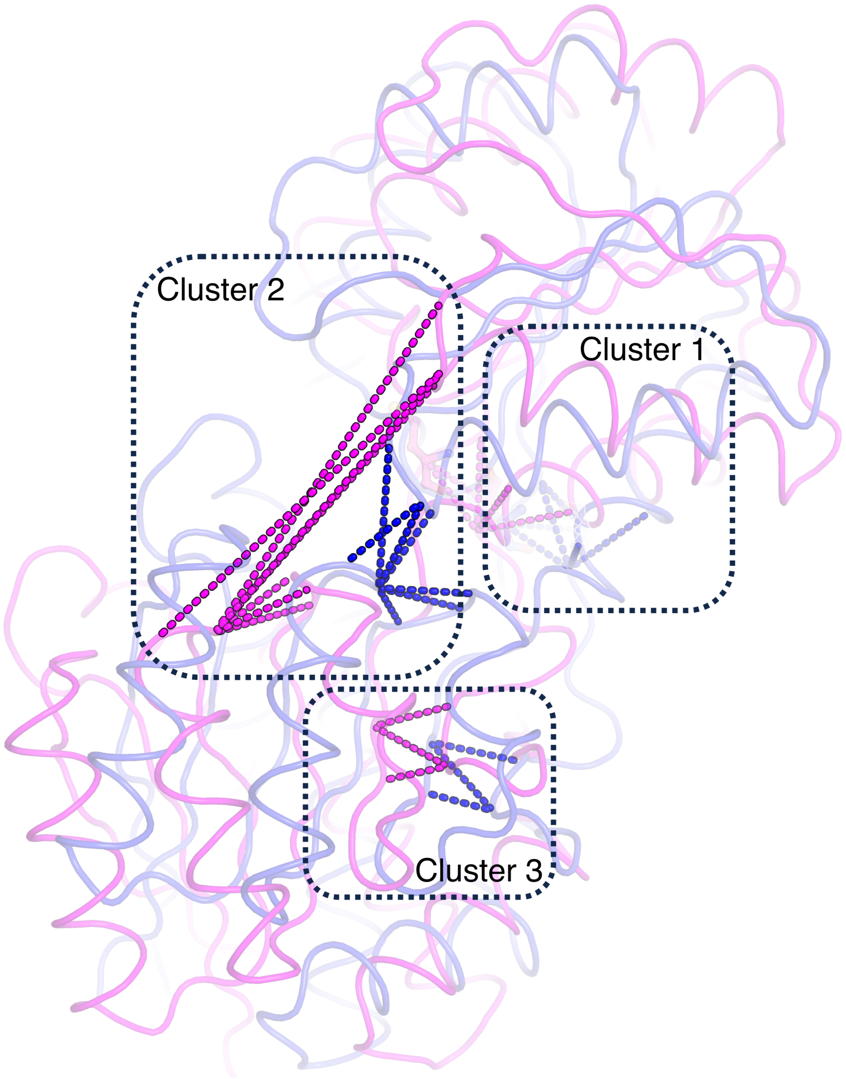


Figure S7: **Illustration of fingerprint contacts for LIVK_ECOLI.** Visualization of the three detected clusters in both closed (PDB: 1USI_A[S3]) and open (PDB: 1USG_A[S3]) crystal structures. The open structure is superimposed onto the closed structure.

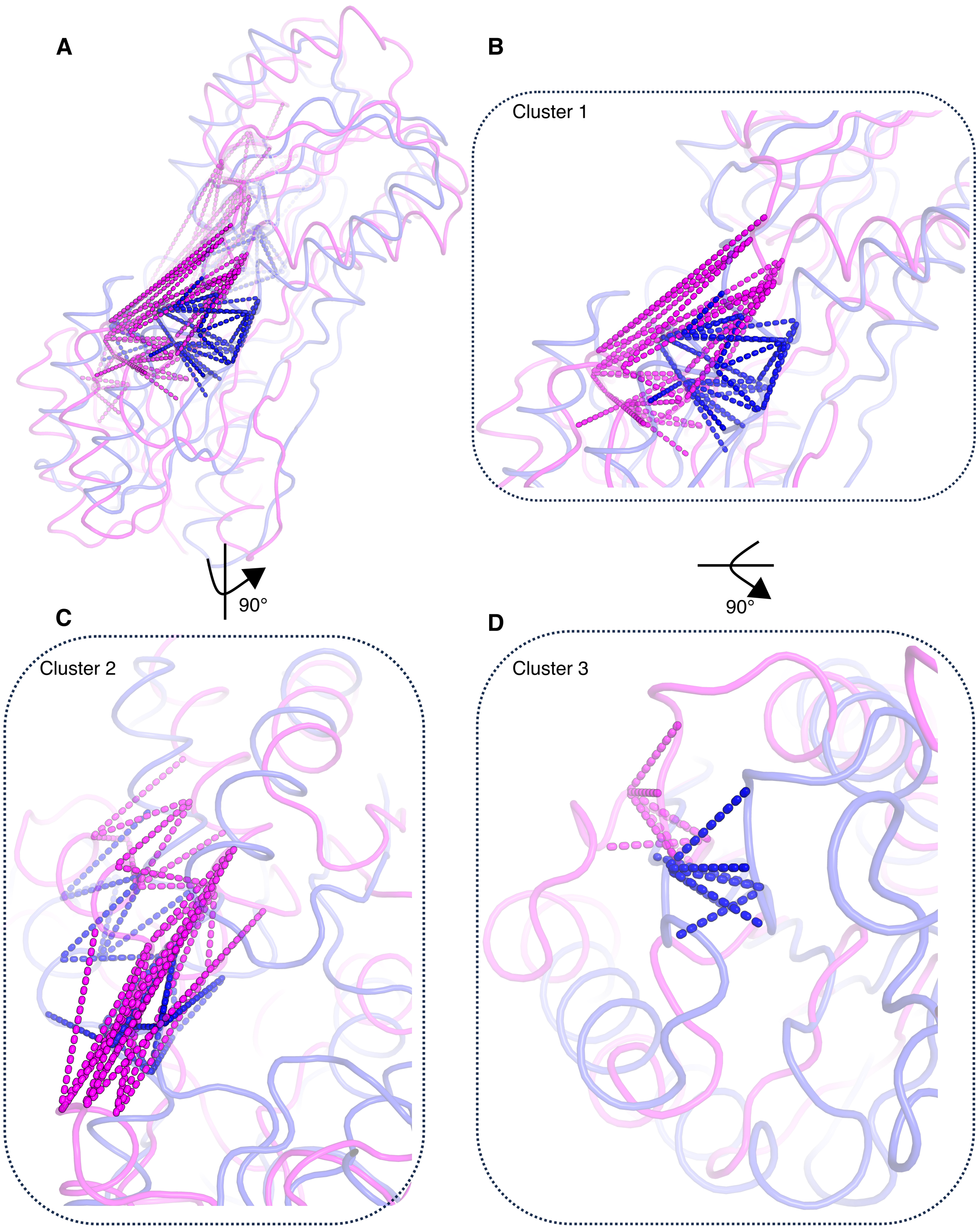


Figure S8: **Illustration of fingerprint contacts for RBSB_ECOLI.** (A) Visualization of the three detected clusters in both closed (PDB: 2DRI_A[S4]) and open (PDB: 1BA2_A[S5]) crystal structures. (B-D) Zoom in view of Cluster 1, 2, and 3. The open structure is superimposed onto the closed structure.

### References


(S1) Müller, C. W.; Schulz, G. E. Structure of the Complex between Adenylate Kinase from Escherichia Coli and the Inhibitor Ap5A Refined at 1.9 Å Resolution: A Model for a Catalytic Transition State. *J. Mol. Biol.* **1992**, *224*, 159–177.

(S2) Müller, C.; Schlauderer, G.; Reinstein, J.; Schulz, G. E. Adenylate Kinase Motions during Catalysis: An Energetic Counterweight Balancing Substrate Binding. *Struc.* **1996**, *4*, 147–156.

(S3) Magnusson, U.; Salopek-Sondi, B.; Luck, L. A.; Mowbray, S. L. X-ray Structures of the Leucine-binding Protein Illustrate Conformational Changes and the Basis of Ligand Specificity. *J. Biol. Chem.* **2004**, *279*, 8747–8752.

(S4) Björkman, A.; Binnie, R. A.; Zhang, H.; Cole, L. B.; Hermodson, M. A.; Mowbray, S. L. Probing Protein-Protein Interactions. The Ribose-Binding Protein in Bacterial Transport and Chemotaxis. *J. Biol. Chem.* **1994**, *269*, 30206–30211.

(S5) Björkman, A. J.; Mowbray, S. L. Multiple Open Forms of Ribose-Binding Protein Trace the Path of its Conformational Change. *J. Mol. Biol.* **1998**, *279*, 651–664.
